# Sorgoleone degradation by sorghum-associated bacteria; an opportunity for enforcing plant growth promotion

**DOI:** 10.1101/2023.05.26.542311

**Authors:** Yasuhiro Oda, Joshua R. Elmore, William C. Nelson, Andrew Wilson, Yuliya Farris, Ritu Shrestha, Citlali Fonseca Garcia, Dean Pettinga, Aaron J. Ogden, Henri Baldino, William G. Alexander, Adam M Deutschbauer, Catalina Vega Hurtado, Jason E. McDermott, Adam M. Guss, Devin Coleman-Derr, Ryan McClure, Caroline S. Harwood, Robert G. Egbert

## Abstract

Metabolite exchange between plant roots and their associated rhizosphere microbiomes underpins plant growth promotion by microbes. *Sorghum bicolor* is a cereal crop that feeds animals and humans and is used for bioethanol production. Its root tips exude large amounts of a lipophilic benzoquinone called sorgoleone. Sorgoleone is an allelochemical that suppresses the growth of competing plant seedlings and is mineralized by microbes in soil. As an avenue to understand how sorghum and its root microbiome may be connected through root exudates, we identified the molecular determinants of microbial sorgoleone degradation and the distribution of this trait among microbes. We isolated and studied from sorghum-associated soils, three bacterial strains classified as *Acinetobacter*, *Burkholderia*, and *Pseudomonas* species that grow with sorgoleone as a sole carbon and energy source. The genomes of these strains were sequenced and subjected to transcriptomic and gene fitness analyses to identify candidate sorgoleone degradation genes. Follow up mutational analysis showed that sorgoleone catabolism is dependent on four contiguous genes that are conserved among the species we sequenced. Phylogenetic analysis of the sorgoleone degradation gene cluster showed that sorgoleone catabolism is enriched in sorghum-associated *Streptomyces* strains. The discovery of bacteria that grow on a compound like sorgoleone that is plant specific and not widely distributed in the environment, provides an opportunity to study how a plant exudate can enforce the development of a rhizosphere specific microbiome for the mutual benefit of plant and microbe.

**Significance:** The grain crop sorghum exudes an herbicidal compound called sorgoleone from its root tips, which inhibits the growth of other plants. We isolated bacteria that grow on sorogleone and identified a cluster of bacterial genes required for sorogleone degradation that can be used as a biomarker for this trait. An approach to improve the production of crops in stressful conditions such as drought, is to encourage their association with plant growth promoting bacteria. Our discovery of sorgoleone degradation genes opens the door to engineering bacteria that receive benefit from sorghum in the form of a plant-specific growth substrate, and in return promote the growth of this crop.

## Introduction

Root rhizosphere bacteria can benefit their host plants (1, 2) by providing nitrogen and phosphorus (3–5), reducing infection from plant pathogens (6), and mitigating various stress conditions, such as drought (7, 8). Most plants exude small organic compounds including organic acids, sugars and amino acids that support the growth of the bacteria in their rhizospheres (9). In addition, plants exude plant-specific secondary metabolites that play a role in plant protection from pathogens and are important for shaping the rhizosphere microbiome (10–12). Given the large number of secondary metabolites produced by plants, there are still relatively few known mechanisms by which they exert their effects on microbes. Use of secondary metabolites as carbon sources for growth is an obvious mechanism to shape microbiome composition but has received relatively little attention.

The secondary metabolite sorgoleone, 2-hydroxy-5-methoxy-3-[(8’*Z*,11’*Z*)-8’,11’,14’- pentadecatriene]-*p*-benzoquinone (Fig. 1), is a major component of exudates from root seedlings of *Sorghum bicolor*, the fifth largest cereal crop worldwide (13, 14). Sorgoleone has drawn substantial attention from the research community and the agriculture industry because of its allelochemical properties (15, 16). It has a variety of impacts on soil ecology and has been used in integrated weed management (17). It can also reduce loss of nitrogen fertilizers in soil by inhibiting biological nitrification (18). Previous work has shown that sorgoleone is slowly mineralized to carbon dioxide in soils and that microbial activities are responsible for this process (19). Sorgoleone also influences the composition and network structure of soil and rhizosphere microbial communities (20).

**Fig. 1.**
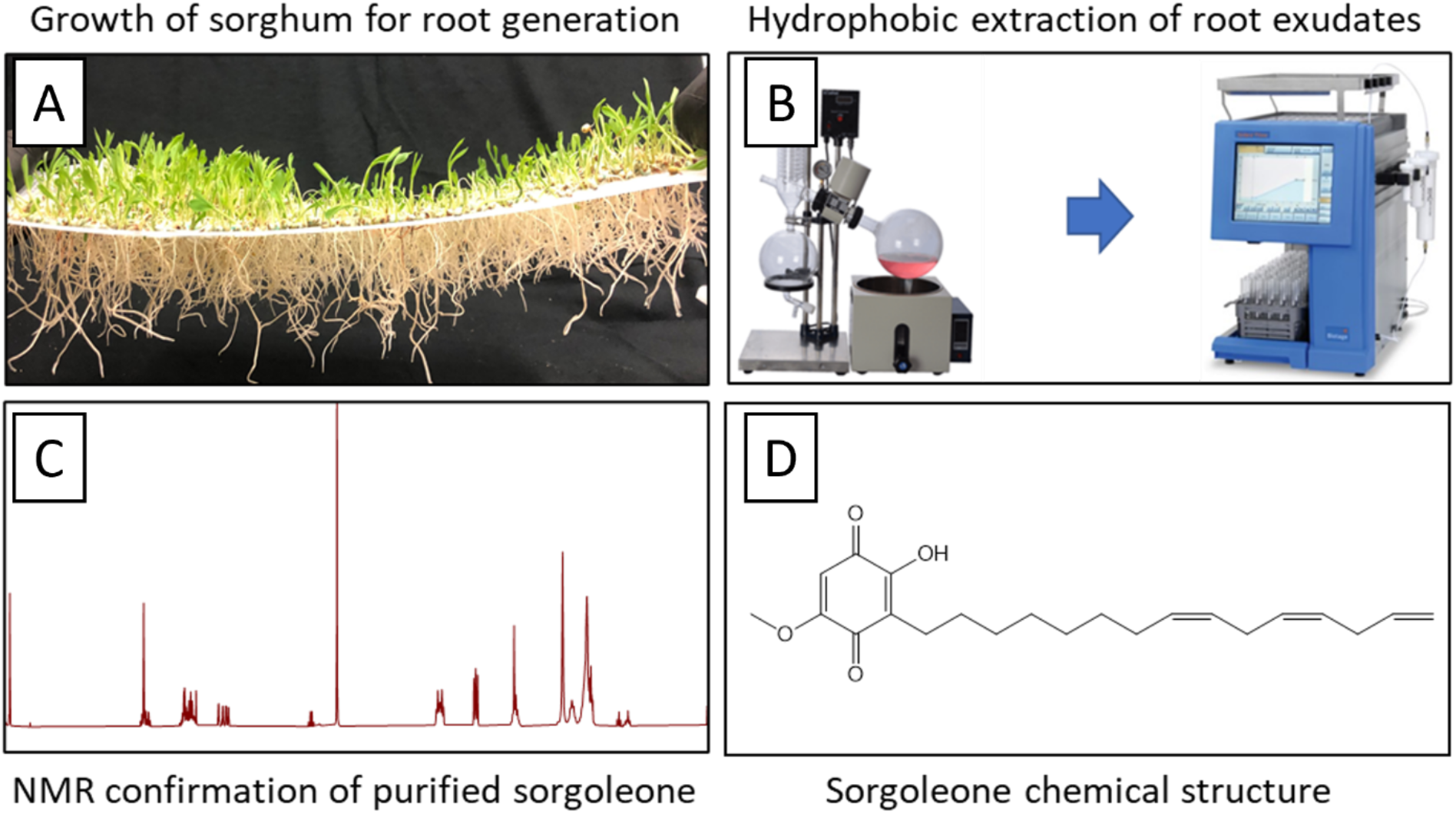
Summary of sorgoleone isolation. (A) Germination of sorghum seeds. (B) Extraction of sorgoleone from germinated sorghum roots using MeOH/dichloromethane and fractionation by thin layer chromatography. (C) Qualification of purified sorgoleone by NMR analysis. (D) Chemical structure of sorgoleone.

Here, we enriched and isolated sorgoleone-degrading bacteria from soil that had been planted with sorghum by using sorgoleone as a sole carbon source. Whole genome sequencing of three sorgoleone-utilizing isolates followed by transcriptome and RB-TnSeq analyses identified genes likely to be involved in sorgoleone degradation. Mutational analysis confirmed that a four gene cluster conserved among our isolates was required for growth on sorgoleone. A phylogenetic survey further revealed that these genes were enriched in *Streptomyces* species associated with sorghum. These results are a first step to determine if sorgoleone might be harnessed to control persistence of plant beneficial bacteria in root rhizospheres.

## Results

### Isolation and sequencing of sorgoleone degrading bacteria

Sorgoleone was purified from sorghum seedlings (14) as summarized in Fig. 1, and used as a sole carbon source to enrich and isolate three sorgoleone-degrading strains from soli collected from a sorghum-growing field site in Kearney, CA, USA. Two strains, SO1 and SO82 grew with 2 mM sorgoleone to a final yield of about 10^9^ CFU/mL. The third strain, SO81, grew to lower yields on the same concentration of sorgoleone (Fig. 2). A combination of whole genome sequencing and Genome Taxonomy Database (GTDB) analysis (https://gtdb.ecogenomic.org/), designated strains, SO1 and SO82, as *Acinetobacter pitii* and *Burkholderia anthina*, respectively (*SI Appendix*, Table S1). Strain SO81 was a novel *Pseudomonas* species. Based upon its ability to use sorgoleone as a carbon source, we named this strain *Pseudomonas sorgoleonovorans* SO81.

**Fig. 2.**
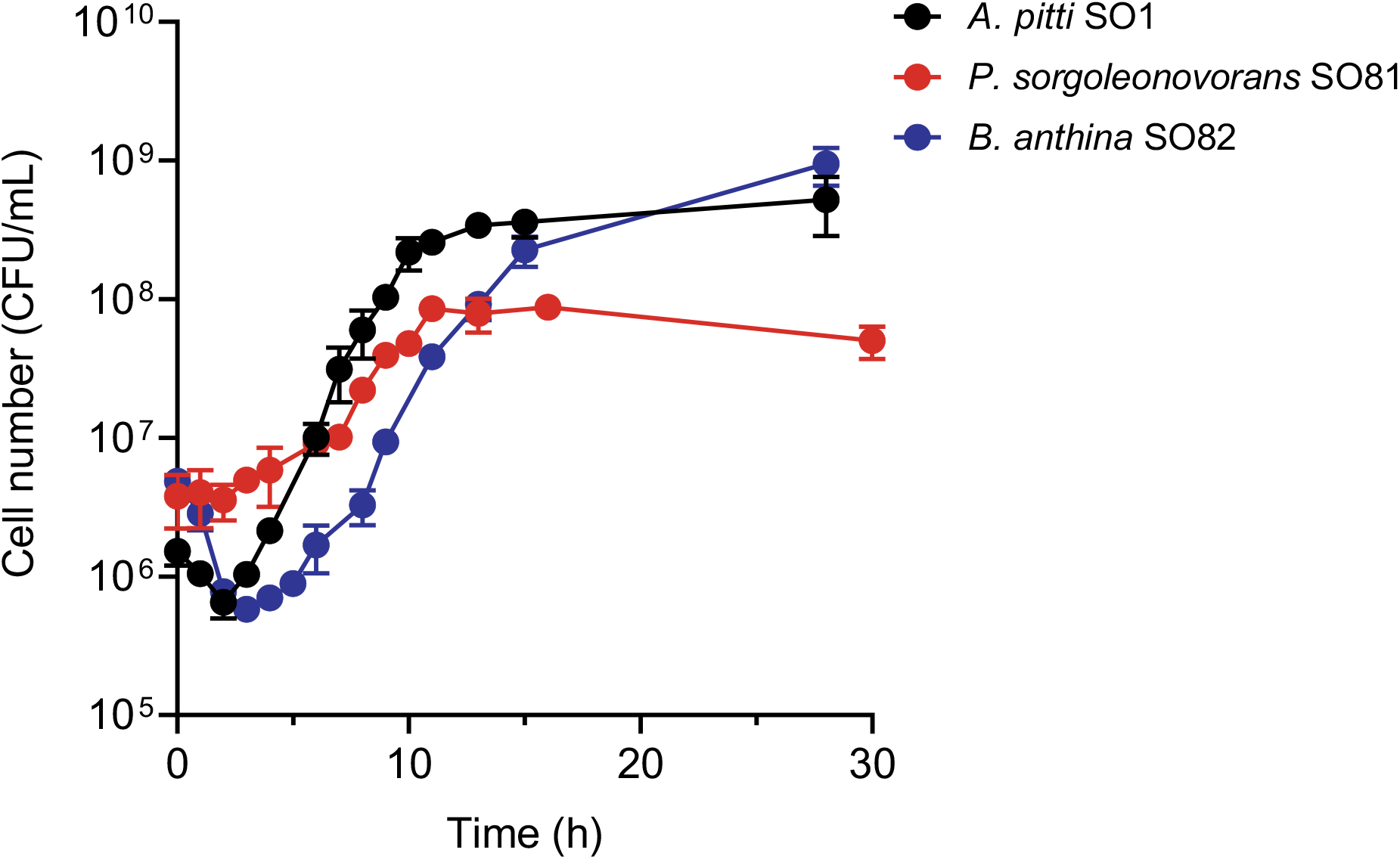
Growth of *A. pitti* SO1, *P. sorgoleonovorans* SO81, and *B. anthina* SO82 on 2 mM sorgoleone as a sole carbon and energy source. Growth as expressed as colony forming units (CFU). The averages of three replicate cultures are plotted with error bars showing the standard deviations.

### Transcriptome analysis to identify genes expressed at high levels during growth with sorgoleone

Sorgoleone is structurally complex compound comprised of a methoxylated and hydroxylated 1,4 benzoquinone with a polyunsaturated aliphatic side chain (Fig. 1). Its degradation is thus likely to require multiple enzymatic steps. To identify sorgoleone degradation genes, we determined the transcriptomes of each strain grown with either sorgoleone or acetate as sole carbon sources (*SI Appendix*, Fig. S1 and Table S2). We found that 228 genes from *A. pitti* SO1 and 156 genes from *B. anthina* SO82 were expressed at ≥8- fold higher levels in cells grown with sorgoleone as compared to acetate. By contrast just 33 genes were expressed at ≥8-fold higher levels in *P. sorgoleonovorans* SO81 grown with sorgoleone as compared to acetate.

The three strains shared seven genes in common that were expressed at high levels during growth with sorgoleone (Table 1). These included four genes predicted to encode a monooxygenase, two α/β hydrolases, and a cytosine deaminase. Each of these genes was expressed at greater than 40-fold higher levels during growth on sorgoleone compared to acetate. Except for *P. sorgoleonovorans* SO81, which has a permease gene (SO81_17470) between its monooxygenase and cytosine deaminase genes, the four genes are adjacent to each other on the genomes. We therefore refer to these four genes as the *srg* (sorgoleone degradation) cluster (Fig. 3). Other genes highly expressed in the three strains during growth on sorgoleone included an alkane 1-monooxygenase, a 2,4-dienoyl-CoA reductase, and an acyl-CoA dehydrogenase that may be associated with fatty acid β-oxidation.

**Fig. 3.**
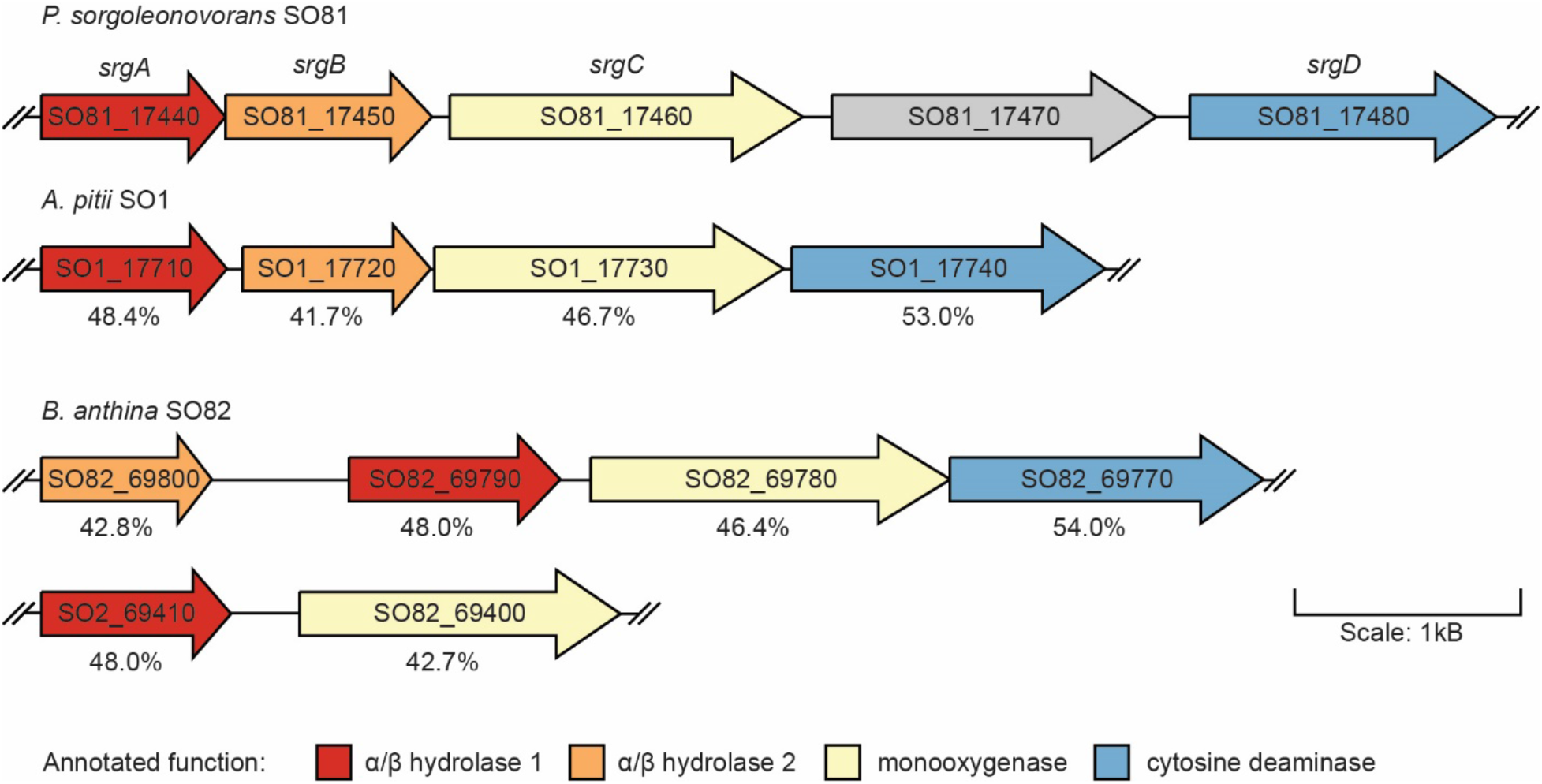
The *srg* cluster for sorgoleone degradation. The color code indicates similar functions among strains. Percentages indicates the protein identity to *P. sorgoleonovorans* SO81 genes.

**Table 1.** Genes highly expressed in strains grown on sorgoleone^a^

|  | *A. pittii* | *P. sorgoleonovorans* | *B. anthina* |
| --- | --- | --- | --- |
| Predicted gene product | SO1 locus | SO81 locus | SO82 locus |
| $\alpha/\beta$ fold family hydrolase | SO1_17710 (467) | SO81_17440 (231) | SO82_69410 (294) |
| $\alpha/\beta$ fold family hydrolase | SO1_17720 (131) | SO81_17450 (165) | SO82_69800 (7.5) |
| Monooxygenase | SO1_17730 (80) | SO81_17460 (199) | SO82_69780 (258) |
| Cytosine deaminase | SO1_17740 (41) | SO81_17480 (96) | SO82_69770 (347) |
| Alkane 1-monooxygenase | SO1_22090 (117) | SO81_11930 (3.4) | SO82_20560 (6.7) |
| 2,4-dienoyl-CoA reductase | SO1_25070 (3.9) | SO81_08370 (4.1) | SO82_24270 (15) |
| Acyl-CoA dehydrogenase | SO1_34600 (2.6) | SO81_19860 (3.6) | SO82_06880 (19) |

<sup>a</sup> Number in parentheses indicates the fold change in expression relative to acetate-grown cells.

Nearly 80% of the genes that were highly expressed in sorgoleone-grown cells for each strain, were either strain-specific or upregulated in just one strain, even if the gene was shared with the other strains. There were species-specific genes adjacent to the *srg* cluster that were highly expressed. For *P. sorgoleonovorans* SO81, several membrane-associated proteins (SO81_17470, SO81_17500 and SO81_17510) and genes associated with β-oxidation processes (SO81_17530 and SO81_17540) were highly expressed, and for *B. anthina* SO82, a quinone oxidoreductase gene (SO82_69480) and a succinate-semialdehyde dehydrogenase gene (SO82_69450) were highly expressed. Genes elsewhere on genomes that were highly expressed in sorgoleone-grown cells in a strain-specific way included genes for benzoate (SO1_16420 to 16480) and 4-hydroxybenzoate (SO1_24810 to 24930) degradation as well as genes for malonic acid degradation (SO1_20050 to 20130) in *A. pitti* SO1. For *P. sorgoleonovorans* SO81, genes likely to be involved in fatty acid degradation (SO81_09470, SO81_12220, SO81_17530, and SO81_17540) and an additional copy of a 2,4-dienoyl-CoA reductase gene (SO81_35370) were expressed at higher levels during growth on sorgoleone. For *B. anthina* SO82, in addition to fatty acid and dicarboxylic acid degradation genes (SO82_36570, SO82_38510, and SO82_42710), three additional copies of genes (SO82_32580, SO82_32670, and SO82_69790) for α/β hydrolase fold enzymes and an additional copy of a 2,4-dienoyl-CoA reductase gene (SO82_43600) were highly expressed. These results suggest that the three strain strains may catabolize sorgoleone to generate different intermediates and downstream products. This may also explain why *P. sorgoleonovorans* SO81 did not grow to as high a yield on sorgoleone as the other two strains we studied.

We found that genes associated with oxidative and other stress responses were more highly expressed in *A. pitti* SO1 cells grown with sorgoleone. These included a universal stress protein gene (SO1_25550), genes involved in trehalose biosynthesis (SO1_12530 and SO1_12540), and a gene cluster containing catalase (SO1_19670 to SO1_19740). In *P. sorgoleonovorans* SO81, genes for a multidrug efflux system (SO81_01900 to 01920 and SO81_30880 to 30900) were upregulated.

### RB-TnSeq analysis to identify candidate genes for sorgoleone degradation

We used the functional genomics-based approach of RB-TnSeq to screen directly for genes involved in sorgoleone catabolism (21). Because of the ease of genetic manipulation of *Pseudomonas* species, we generated an RB-TnSeq library in *P. sorgoleonovorans* SO81. The library had 367,775 unique barcoded transposon insertion mutants with insertions in 3,924 of the 4,649 predicted genes (84%), with an average coverage of 60.3 transposon insertion mutants per gene. Using this library, we performed fitness assays that evaluated the relative growth of mutants in the library on sorgoleone and a series of other carbon sources (acetate, citrate, glucose, or octanoate). Genes were considered important for robust growth under a given condition if their fitness score was ≤-1.8 (strong negative fitness) (*SI Appendix*, Table S3).

We identified 14 genes that had strong negative fitness scores only when the library was grown with sorgoleone. Of these, eight genes were induced by sorgoleone in our transcriptome analysis (Fig. 4). Included in this set were the *srg* cluster (SO81_17440 to 17480) and two genes adjacent to the *srg* cluster. Our RB-TnSeq experiments identified several genes involved in β-oxidation (SO81_19340, SO81_19860, SO81_39590, SO81_42070, and SO81_42080) that conferred substantially reduced fitness during growth on either sorgoleone or ocanoate when disrupted (Fig. 4). We also identified several genes potentially involved in the transport of sorgoleone and/or intermediates of its degradation. Among these were SO81_12750 to 12770 (putative ABC transporter), SO81_18010 (murein hydrolase transporter LrgA), and SO81_30590 (channel protein TolC). SO81_17430, which encodes a predicted transcription factor (*SI Appendix*, Table S3), had a positive fitness score, suggesting that it negatively regulates transcription of the *srg* cluster.

**Fig. 4.**
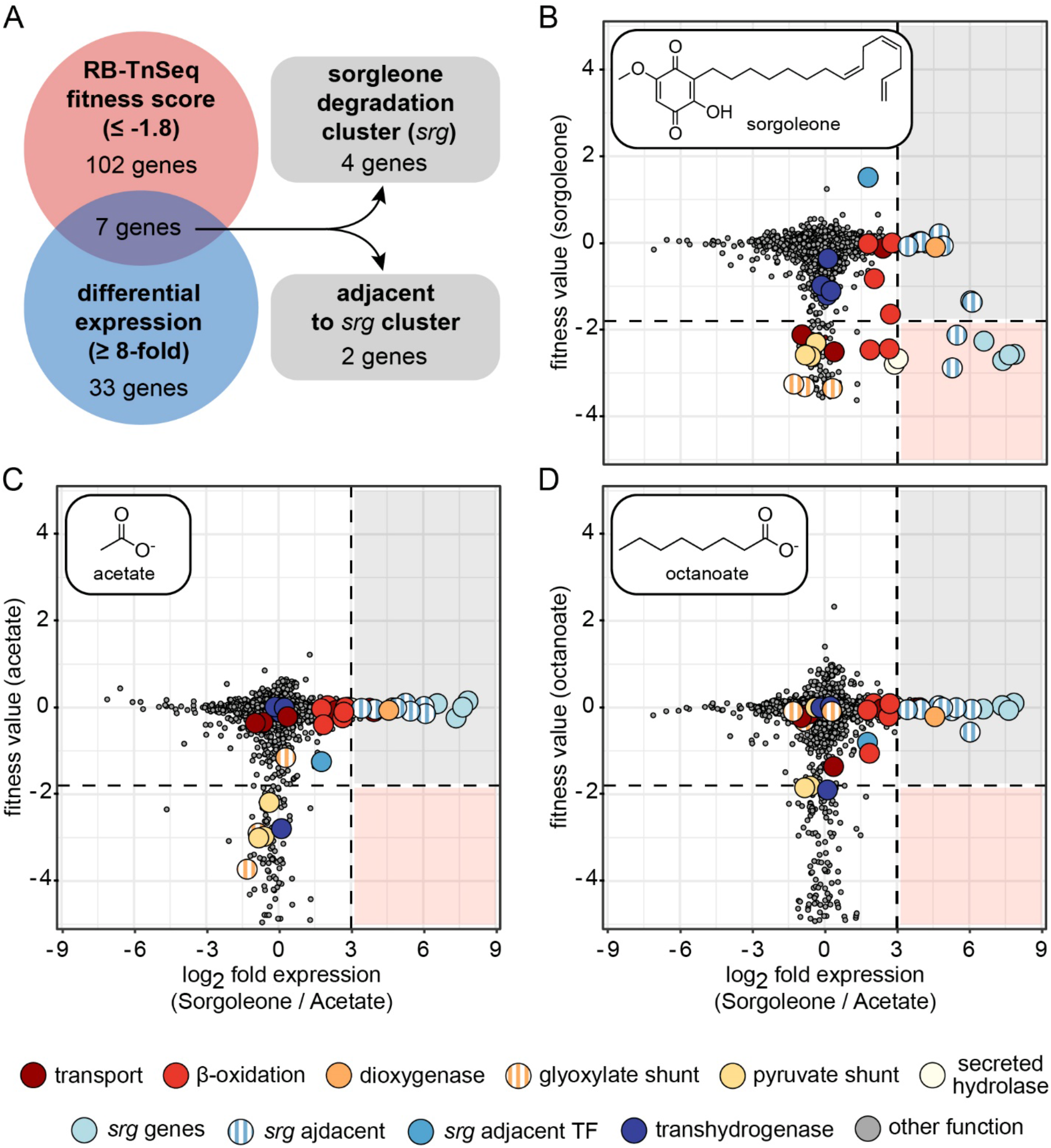
Summary of RB-TnSeq and RNA-Seq data from *P. sorgoleonovorans* SO81. (A) Venn diagram showing overlap between genes with substantial fitness values when *P. sorgoleonovorans* SO81 is grown on sorgoleone and genes that are more highly expressed when *P. sorgoleonovorans* SO81 is cultivated in sorgoleone versus acetate as carbon sources. Plots comparing differential expression (x-axis) versus mean RB-TnSeq fitness values on the y-axis. RB-TnSeq values displayed are from cultures grown on either (B) sorgoleone, (C) acetate, or (D) octanoate. Positive and negative differential expression values indicate higher expression during growth on sorgoleone and acetate as carbon sources, respectively. Dots indicate transporters (dark red), β-oxidation enzymes (red), dioxygenase (orange), glyoxylate shunt enzymes (orange stripes), pyruvate shunt enzymes (yellow), a putative secreted hydrolase (light yellow), the *srg* cluster (light blue), genes adjacent to the *srg* cluster (light blue stripes), transcription factor adjacent to the *srg* cluster (blue), NAD(P)H transhydrogenase (dark blue), and all other genes (gray). Values represent the mean of three to four biological replicates of RB-TnSeq analysis and four biological replicates of RNA-Seq differential expression analysis. Genes lacking fitness or differential expression value are not displayed.

### The *srg* cluster is essential for sorgoleone degradation

To confirm that the *srg* cluster is essential for sorgoleone degradation, we constructed a *P. sorgoleonovorans* SO81 *srg* deletion mutant (SO81Δ*17440-17460*). Both the wild type and the deletion mutant grew well on glucose, but the mutant strain failed to grow on sorgoleone. Integration of a single copy of the *srg* cluster (*SO81_17440-17460*) into the *attTn7* site of SO81Δ*17440-17460* complemented this phenotype (*SI Appendix*, Fig. S2).

### The *srg* cluster as a biomarker for sorgoleone catabolism

Our observation that the *srg* cluster is conserved among several species of our soil isolates prompted an investigation of the distribution of this gene cluster across sequenced organisms and its suitability as a genetic marker for sorgoleone catabolism. Our approach involved cluster construction of Snekmer protein family models (22) for the *srg* genes using sequences from our three isolates and data from the UniProt Reference Proteome. We used the *srg* protein models to search the GFOBAP collection of 3,837 genomes derived from soil- and plant-associated environments (23), and identified hits for the complete cluster in one *Acinetobacter* and 21 Actinobacteria, most of which are *Streptomyces* sp. (*SI Appendix*, Fig. S3).

To investigate the prevalence of the *srg* cluster among *Streptomyces* isolates associated with different plants we analyzed publicly available whole genome sequences of 44 *Streptomyces* strains isolated from sorghum-cultivated fields and 30 *Streptomyces* strains isolated from soils near poplar trees. We found a significant enrichment of the *srg* cluster among the strains associated with sorghum (17/44) over poplar (4/30) (two-proportion z-test, p=0.012) (Fig. 5). To assess whether the presence of the *srg* cluster is required for *Streptomyces* to grow on sorgoleone, we selected a taxonomically diverse collection of sorghum-associated *Streptomyces* strains, including strains with and without the *srg* cluster, and tested for growth in minimal medium containing sorgoleone as the sole carbon source. We found that only the strains encoding the *srg* cluster were able to grow on sorgoleone (Fig. 5). An ortholog profiling analysis of *Streptomyces* strains revealed general conservation of the gene neighborhood of the *srg* cluster with several strains encoding a TetR family transcriptional regulator adjacent to the *srg* cluster (*SI Appendix*, Fig. S4). These results suggest that the *srg* cluster provides the critical steps to break down sorgoleone in *Streptomyces* and can be used as a biomarker for sorgoleone catabolism.

**Fig. 5.**
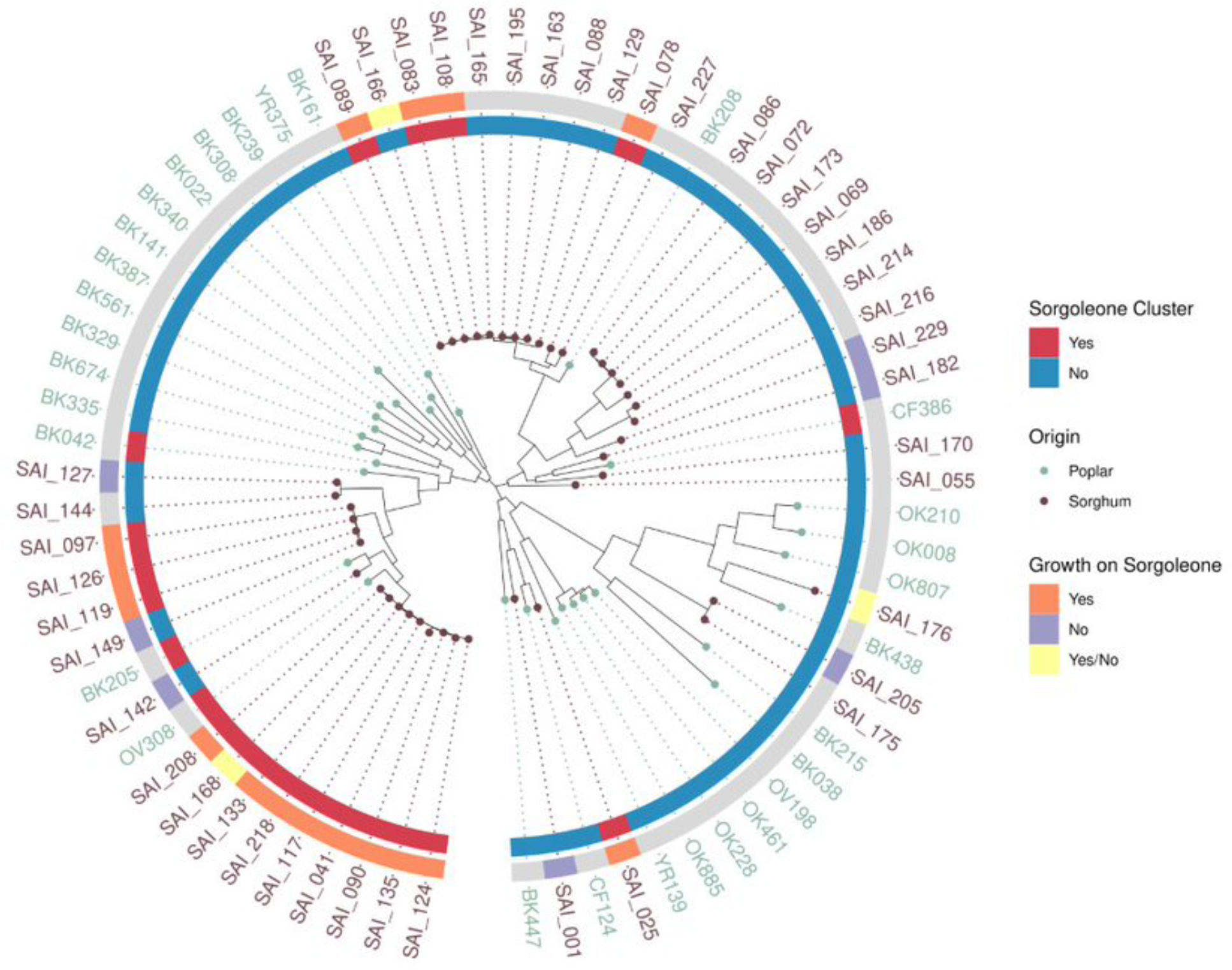
Phylogenetic analysis and growth of selected *Streptomyces* strains on sorgoleone.

## Discussion

Microbial rhizosphere communities are complex. Plants can have both positive and negative effects on their associated root microbes and there is also a complex web of antagonistic and cooperative interactions among the microbial members of the rhizosphere (12, 24). This is against a backdrop of fluctuating soil conditions, particularly soil moisture conditions. Evidence suggests that while many kinds of plants share a core microbiome, plant-specific exudates play a role shaping plant specific microbiomes that include additional strains and species (9).

Here we purified sorgoleone from sorghum seedlings using contemporary techniques and used it to isolate bacteria that grow with sorgoleone as a sole carbon and energy source. This led to the discovery of four contiguous *sgr* genes required for growth with sorgoleone as a carbon source. A next step will be to elucidate the enzymatic activities of the Sgr proteins. The Integrated Microbial Genome (IMG) database (https://img.jgi.doe.gov/) sometimes has more specific gene annotations than the pipeline that we used to annotate the genomes reported here. Using IMG annotations for the *srg* genes in *P. sorgoleonovorans* SO81, we can hypothesize a possible route of sorgoleone degradation. The predicted proteins of SO81_17440 and SO81_17450 are most closely related to pimeloyl-ACP methylester carboxylesterase, which is involved in biotin synthesis (25). We hypothesize that these genes could catalyze demethoxylation of sorgoleone in *P. sorgoleonovorans* SO81. SO81_17460 is related to 2- polyprenyl-6-methoxy phenol hydroxylase-like FAD dependent oxidoreductase, which is required for ubiquinone synthesis (26). This flavoprotein monooxygenase enzyme could catalyze oxidative rearrangement of the benzoquinone ring resulting in a lactone (a Baeyer-Villiger-type reaction) or another intermediate which then easily undergoes hydrolytic or even spontaneous ring cleavage. The cytosine deaminase predicted to be encoded by the fourth gene (SO81_17480) has a general role as a metal-dependent hydrolase and could play a role in the cleavage of a lactone. Other genes shared among the three strains (Table 1) encode enzymes that are known to catalyze fatty acid degradation, and these could be important for degradation of the unsaturated fatty acid tail on sorgoleone. The genes in the *srg* cluster are related to co-factor biosynthesis and it may be that this is their evolutionary origin.

Our observation that the *srg* cluster is enriched in sorghum-associated *Streptomyces* isolates suggests that this group may have a competitive growth advantage over other microbes in the sorghum rhizosphere. A recent study observed that *Streptomyces* can reach comparatively high abundances in the sorghum rhizosphere (27). And *Streptomyces* have been shown to be active in promoting the growth of many crop plants (28) including sorghum (29, 30). We suggest that it may be possible to take advantage of sorgoleone exudation by sorghum to enforce persistence of sorghum-utilizing bacteria that can promote the growth of this important food and bioenergy crop. To do this it will be important to understand a complete picture of sorgoleone production at each point in the life cycle of sorghum and to fully elucidate how sorgoleone is degraded.

## Materials and Methods

### Purification of sorgoleone from sorghum

To generate sorgoleone extracts, *Sorghum bicolor* SX-19 (hereafter referred to as SX-19) seedlings were grown in bulk using a devised aeroponic system. SX-19 seeds (Helena Agri-Enterprises) were first soaked in sterile deionized water for at least 3 h. After soaking, the SX-19 seeds were spread across a plastic mesh canvas (85 g of dry seeds per mesh canvas) which was placed on risers in a shallow plastic tray. The seeds were then covered with damp paper towels, the trays were covered with aluminum foil, and placed either in a growth chamber at 20 °C or on a benchtop at room temperature. All plastic was sterilized by autoclave prior to planting, and the seeds were handled in a laminar flow hood to minimize microbial contamination. The trays were watered every 24 to 48 h by re-wetting the paper towels with sterile water. Post germination, the SX-19 roots grew through the plastic mesh into the headspace between the mesh and the bottom of the tray. After 5 to 10 days of growth, SX-19 roots were harvested by lifting the plastic screen and excising the roots that had grown through the mesh. Excised roots were placed in a glass beaker and covered with HPLC grade chloroform for 2 min to extract any hydrophobic root exudates. The extract was decanted into a liter glass bottle, capped, and stored for up to one week until purification. The chloroform treated roots were dried and weighed. This method yielded about 0.015 g crude extract per g of dry root mass on average (n=18 ± 0.01).

Sorgoleone was purified from the crude extract using a Biotage Isolera flash chromatography system. Crude extract was dissolved in chloroform or dichloromethane and loaded onto pre-packed silica gel columns equilibrated with 1% MeOH/dichloromethane. Pure sorgoleone was eluted by ramping from 1 to 5% MeOH/dichloromethane. Fractions of product were combined and dried under reduced pressure to afford pure sorgoleone as a bright orange solid. The average purified sorgoleone yield was 0.0067 g per g of dry root mass (n=18 ± 0.0024).

Purified sorgoleone was characterized using nuclear magnetic resonance (NMR) spectroscopy and low-resolution mass spectrometry (LRMS), and compared to the published data (31). ^1^H and ^13^C NMR spectra were acquired in CDCl_3_ (Cambridge Isotopes, Tewksbury, MA) at 25 °C on a Bruker 400 MHz Avance III spectrometer equipped with a 5 mm BBFO SmartProbe. All chemical shifts are reported in the standard notation of parts per million using tetramethylsilane as an internal reference or the peak of the residual proton or carbon signal of CDCl_3_ (^1^H NMR δ 7.26 and ^13^C NMR δ 77.36 ppm) as an internal reference. LRMS-ESI was performed using a Finnigan LTQ mass spectrometer (Thermo Electron Corporation). Purified sorgoleone was dissolved in 100% DMSO to a final concentration of 200 mM for use in microbial cultivation.

### Bacterial growth conditions

Bacteria were generally grown in MME, MMV (MME with 1x vitamin solution), or M9 medium with variable carbon sources. MME medium contains 9.1 mM K_2_HPO_4_, 20 mM MOPS, 4.3 mM NaCl, 9.3 mM NH_4_Cl, 0.41 mM MgSO_4_, 68 µM CaCl_2_, 1x MME trace minerals, and final pH was adjusted to 7.0 with KOH. The 1000x MME trace mineral stock solution contains 1 mL concentrated HCl, 0.5 g Na_4_EDTA, 2 g FeCl_3_, 0.05 g each H_3_BO_3_, ZnCl_2_, CuCl_2_×2H_2_O, MnCl_2_×4H_2_O, (NH_4_)_2_MoO_4_, CoCl_2_×6H_2_O, NiCl_2_×6H_2_O per liter. The 200x vitamin solution contains 10 mg each of niacin, pantothenate, lipoic acid, *p*-aminobenzoic acid, thiamine (B_1_), riboflavin (B_2_), pyridoxine (B_6_), and cobalamin (B_12_) and 4 mg each of biotin and folic acid per liter. Solid medium was prepared by addition of 15 g of Bacto Agar (BD) per liter. Routine cultivation of the sorgoleone-utilizing isolates was performed using Difco R2A (BD), Difco SOB medium (BD), or Difco LB, Miller (BD).

### Isolation of sorgoleone-utilizing bacteria by enrichment technique

Rhizosphere soil was collected from the Kearney Agricultural Research and Extension (KARE) Center in Parlier, California, where *Sorghum bicolor* has been cultivated. MME medium supplemented with 2 mM sorgoleone (5 mL in glass test tubes) was used to enrich bacteria that could use sorgoleone as a sole carbon source from 100 mg of this rhizosphere soil. After 96h incubation at 25 °C, the initial enrichment culture was diluted 1:100 into fresh MME supplemented with vitamins and containing 2 mM sorgoleone. After 72 h incubation at 25 °C, 100 µL of this culture was spread onto MME agar plate containing 2 mM sorgoleone and incubated for 96 h at 25°C. Well separated single colonies were picked and streaked onto R2A agar plates for single colony isolation. Selected isolates were inoculated into MME containing 1 mM sorgoleone. Isolates grown in this medium were serially passaged (1:100 dilution) six times into fresh MME containing 1 mM sorgoleone. These cultures were spread onto R2A, and single colonies were inoculated into R2A for growth prior to storage as glycerol stocks at -80 °C. The 16S rRNA sequences of the isolates were determined by the Sanger sequencing (GENEWIZ).

### Whole genome sequencing and analysis

Whole genomes of isolates were sequenced by CD Genomics (Shirley, NY, USA) and their taxonomies were assigned using Genome Taxonomy Database-Tk v1.6.0 with the classify_wf settings and reference data version r202 (32).

Functional annotation was assigned using the DFAST pipeline (v 1.2.14) (33) supplemented with RAST (34), KOfam (35) (2021-06-01 release), and searches against TIGRfam (v15.0) (36) and Pfam (v32.0) (37) using hmmer 3.3 (38). Orthology between genomes was determined using bespoke perl code which integrates reciprocal best hit information, TIGRfam equivalog family membership and conserved synteny.

### Transcriptome analysis of sorgoleone degrading isolates

Sorgoleone-utilizing strains were revived from -80 °C glycerol stocks by cultivation in 5 mL R2A liquid medium at 30 °C. Cultures were centrifuged at 3000 x g and pellets were washed twice with 5 mL MME lacking carbon source to remove residual R2A medium. Starter cultures were inoculated at a 1:100 dilution of the washed cells into 5 mL of MME supplemented with either 2 mM sorgoleone or 30 mM acetate and grown to stationary phase. Final cultures were inoculated at a 1:100 dilution into 50 mL of MME supplemented with either 2 mM sorgoleone or 30 mM acetate and incubated at 30 °C until they reached mid-log-phase (OD_600_ of 0.4 to 0.5 for *A. pitti* SO1 and *B. anthina* SO82, and 0.15 to 0.25 for *P. sorgoleonovorans* SO81). Cultures were centrifuged at 12,000 x g for 2 min at 4 °C, and cell pellets were flash frozen in liquid nitrogen and stored at -80 °C. RNA isolation, rRNA depletion, and high-throughput RNA sequencing with the Illumina sequencing platform was performed by CD Genomics (39, 40).

The transcriptomics data were indexed and quantified by kallisto (41) as per instructions in the manual. The resulting read counts were processed using a bespoke Python code that ran DEseq2 (42) to normalize read counts and to calculate the fold change value between sorgoleone-grown and acetate-grown cells with an adjusted *p*-value calculated for statistical analysis for each gene. Fold differences between expression values of a given gene from cells grown under the two conditions were considered significant if the genes had values with a base mean of ≥100 and an adjusted *p-value* of <0.05.

### RB-TnSeq analysis

A transposon mutant library of *P. sorgoleonovorans* SO81 was constructed by conjugation with *E. coli* WM3064 harboring pHLL250 mariner transposon vector library (strain AMD290) (43). Approximately equal numbers of mid-log-phase grown cells of *P. sorgoleonovorans* SO81 and AMD290 were mixed in SOB containing 300 µM of 2,6- diaminopimelic acid (DAP), incubated at 30 °C on 0.22 µm nitrocellulose filters (Millipore) overlaid on SOB agar plates containing DAP. After 6.5 h, cells were removed from the filters, resuspended in SOB containing 25% glycerol, and stored at -80 °C as a master conjugation mixture. Aliquots of conjugation mixture were plated on SOB agar plates with 50 µg/mL kanamycin (Km) to select for mutants. After 2 days of growth at 30 °C, colonies were scraped from plates and resuspended in SOB. The mutant library was diluted to a starting OD_600_ of 0.25 in 50 mL of SOB with 50 µg/mL Km, cultivated at 30 °C to a final OD_600_ of 1.25, made multiple - 80 °C freezer stocks following addition of glycerol to a final volume of 25%. Equal volumes of 15 independently prepared mutant libraries were pooled to form the final *P. sorgoleonovorans* SO81 mutant library.

We mapped transposon insertion sites and linked these insertions to their associated DNA barcodes using a two-step PCR based approach that selectively amplifies the transposon insertion junctions and adds Illumina adapter sequences (44). For the *Pseudomonas*_S08_1_ML2 library, we prepared three independent Illumina sequencing libraries and sequenced these using paired end sequencing (2 x 150) on an Illumina HiSeq machine (Novogene). The total number of paired reads sequenced across the 3 libraries were 25.4 million, 65.7 million, and 79.3 million. From these data, we were able to confidently map 367,775 unique insertions (with at least 10 reads of support for each) using previously described criteria (21). We performed barcode sequencing (BarSeq) (45) with both P1 and P2 oligos indexed to minimize the impact of misassigned indexes in Illumina HiSeq4000 runs (https://www.biorxiv.org/content/10.1101/125724v1). We followed established methods for calculating strain and gene fitness scores, along with a t-like statistic that describes the confidence in the gene fitness score (21). All the RB-TnSeq software used in this study is available at https://bitbucket.org/berkeleylab/feba/src/master/.

### RB-TnSeq fitness assays

RB-TnSeq assays were performed as previously described (21). Briefly, the *P. sorgoleonovorans* SO81 transposon libraries were thawed at room temperature and grown in 25 mL LB at 37 °C with shaking at 200 rpm until the OD_600_ reached 0.5. Cultures were centrifuged, washed twice with MME, and resuspended to an OD_600_ of 2.0. Aliquots of washed cells were centrifuged, and the resulting cell pellets were stored at -80 °C for later amplicon sequencing and barcode quantification. These aliquots served as the T=0 samples for RB-TnSeq. The remaining washed cells were diluted to a starting OD_600_ of 0.02 in 3 mL MME supplemented with 2 mM sorgoleone in glass test tubes or 1.2 mL MME supplemented with all other carbon sources (30 mM acetate, 10 mM citrate, 10 mM glucose, or 7.5 mM octanoate) in 96-well deep well plates. Glass test tubes and 96-well deep well plates were incubated at 30 °C with shaking at 200 rpm and 1200 rpm, respectively, until stationary phase was reached for each carbon source (typically 24-36 hours). Following completion of growth, cells were harvested for amplicon sequencing and barcode quantification.

### Deletion of the gene cluster SO81_17440 to SO81_17460 in P. sorgoleonovorans SO81

A deletion fragment was synthesized by Integrated DNA Technologies (Coralville, Iowa), cloned into mobilizable suicide vector pSL15A (46) and transformed into *E. coli* NEB 10-beta (New England Bio Labs). The sequence-verified deletion construct was transformed into *E. coli* S17-1, and further mobilized into *P. sorgoleonovorans* SO81 by conjugation on SOB agar plates. Single recombinant conjugants were first selected on M9 plus 10 mM succinate plates containing 20 μg/mL tetracycline. Tetracycline resistant colonies were further plated onto Difco LB agar, Lennox (BD) containing 10% sucrose. Sucrose resistant and tetracycline sensitive colonies were screened by colony PCR and sequencing to validate the expected chromosomal deletion of the gene cluster. To complement the SO81Δ*17440-17460* mutant, a DNA fragment spanning from *SO81_17440* to *SO81_17460* plus the 250 bp upstream that contains a putative transcription start was PCR amplified and cloned into pUC18-mini-Tn7T-Gm (47). The construct was then transformed into *E. coli* NEB 10-beta (New England Bio Labs) for sequence verification. The sequence-verified complementing construct was transformed into *E. coli* S17-1 and this strain was used in a three-strain mating on SOB plates that also included the SO81Δ*17440-17460* mutant and *E. coli* S17-1 carrying the integration plasmid pTNS3. Exconjugants were selected on M9 plates supplemented with 10 mM succinate and 5 μg/mL gentamicin. As a negative control, the empty vector pUC18-mini-Tn7T-Gm was used.

### Identification of sorgoleone catabolism pathways by Snekmer analysis

To examine the distribution of the sorgoleone degradation genes, proteome datasets were searched with kmer composition profile models using the conserved Srg protein sequences. Training sets for each of the genes were built that included the genes in the *srg* cluster from *A. pitti* SO1, *P. sorgoleonovorans* SO81 and *B. anthina* SO82, as well as genes from 25 organisms in the UniProt Reference Proteome (v 2021_03) that shared both sequence similarity and gene synteny. Protein family models were built using Snekmer, a tool that leverages alternate encoding of proteins to account for amino acid substitutions and k-mer based similarity calculation (22). The kmer length and alphabet parameters were optimized using cross-validation of the training set. In brief, we examined the performance of the models using kmer lengths of 4, 8, 12, and 16 and using six different amino acid reduction alphabets including the original sequence (*i.e.*, no recoding). The best performance across the families was achieved with a kmer length of 8 and the MIQS alphabet, which groups amino acids by observed substitution rates (48). A genome was considered positive for sorgoleone genes if it had positive predictions for all three of the gene families that were found in the same genomic region either continuous or separated by no more than three intervening genes to allow more generous detection of the set of functions.

### Phylogenetic analysis of poplar- and sorghum-derived *Streptomyces* isolates

Genomes of *Streptomyces* isolates derived from poplar (n=30) and sorghum (n=43) were used to build a phylogenetic tree with *GToTree* (49). Each genome was searched for 73 single copy genes across bacterial tree of life which were used to build the tree. Briefly, target genes were identified with *HMMER3* v3.2.2 (50), individually aligned with *muscle* v5.1 (51), trimmed with *trimal* v1.4.rev15 (52), and concatenated prior to phylogenetic estimation with *FastTree2* v2.1.11 (53). The resultant tree was plotted with *Ggtree* (54) and annotated with the presence or absence of the *srg* cluster as determined by Snekmer (22) and the capacity to grow on sorgoleone as a sole carbon source.

### Cultivation of sorghum-derived *Streptomyces* on sorgoleone

To validate the ability to metabolize sorgoleone by organisms identified by the Snekmer analysis, we performed growth assays of 27 *Streptomyces* strains using sorgoleone as the sole carbon source. Strains were grown on Tryptic Soy Agar plates at 30 °C for 3 days. Three colonies of similar size were inoculated into glass tubes containing M9 supplemented with 4 mM sorgoleone and incubated for 7 days at 30 °C with shaking at 250 rpm. Additional cultures were made by inoculating each strain in M9 media containing 2 mM glucose (positive control) or in M9 media without carbon source (negative control). For each strain, three biological replicates were carried out. Due to the filamentous morphology of *Streptomyces*, bacterial growth was measured by cell biomass as following: bacterial cells were harvested by centrifugation for 10 min at 12,000 rpm and stored at -20 °C. Frozen pellets were ground in a homogenizer with steel beads for 1 min at 25 Hz. Additionally, cells were hydrolyzed with 1 M NaOH overnight at 4 °C. Bradford reactions were performed in 96-well plates (U-shape Greiner bio-one microplates) with 10 µl of each sample and 200 µl of Coomassie (Bradford) Protein Assay Kit (Thermo Scientific) and incubated for 10 min at room temperature. Absorbance was defeminated at 595 nm using a TECAN infinite M Nano plate reader.

## Data availability

Whole genome sequence data of sorgoleone utilizing strains of *A. pitti* SO1 (GenBank accession CP125227.1), *P. sorgoleonovorans* SO81 (GenBank accession CP126126.1), and *B. anthina* SO82 (GenBank accession pending) have been deposited at GenBank (https://www.ncbi.nlm.nih.gov/genbank/) under the BioProject accession PRJNA878512. Corresponding MIGS.ba.plant-associated.5.0 compliant metadata standards have been provided under BioSample accessions: SAMN31055411 (*A. pitti* SO1), SAMN31055412 (*P. sorgoleonovorans* SO81), and SAMN31055413 (*B. anthina* SO82). CD Genomics PacBio (https://www.cd-genomics.com/PacBio-SMRT-Sequencing.html) long-read raw sequence data have been submitted to the Sequence Read Archive (SRA) (https://www.ncbi.nlm.nih.gov/sra) under the following accessions: SRX17787341 (*A. pitti* SO1), SRX17787342 (*P. sorgoleonovorans* SO81), and SRX17787343 (*B. anthina* SO82). High-throughput RNA-seq data have been deposited (GEO accessions pending) at the NCBI Gene Expression Omnibus (GEO) (https://www.ncbi.nlm.nih.gov/gds/). Primary RNA-seq data submissions contain MINSEQE metadata standards, processed gene expression profile analysis, and raw data files located at SRA (accessions pending). The locus tags of *A. pitti* SO1 (IMG Submission ID 282913), *P. sorgoleonovorans* SO81 (IMG Submission ID 273153), and *B. anthina* SO82 (IMG Submission ID 273157) deposited to the IMG database were used throughout this manuscript, and lookup table of the IMG and the GenBank locus tags can be found in *SI Appendix*, Table S4.

A comprehensive collection of all integrated high-throughput omics data and process method metadata are accessible from the PNNL DataHub (https://data.pnnl.gov/about) institutional repository under the following PerCon SFA project page digital object identifier https://doi.org/10.25584/1969551. PerCon SFA data download DOI contents are structured for DOE compliance leveraging reporting guidelines provided by Genomic Standards Consortium community standard initiatives and stakeholder policies supporting FAIR data principles (https://doi.org/10.25504/FAIRsharing.19ne3m). Data package contents reported here are the first version and contain both primary and secondary Persistence Control of Engineered Functions in Complex Soil Microbiomes (PerCon SFA) project deliverables and subsequent digital data objects. Updated versions of the data reported here are accessible from the drop-down options located under the main PNNL DataHub project page DOI (listed above).

## Acknowledgments

We thank Morgan Price for running the BarSeq analysis and uploading the data to the fitness browser; Susane Fetzner and Simon Ernst for helpful discussion. This research was supported by the Secure Biosystems Design Program, US Department of Energy, Office of Science, Biological and Environmental Research, as part of the PNNL Persistence Control (PerCon) Science Focus Area. Pacific Northwest National Laboratory is operated by Battle for the US Department of Energy under Contract DE-AC05-76RL01830.

## Supporting Information

**SI Appendix, Table S1.**
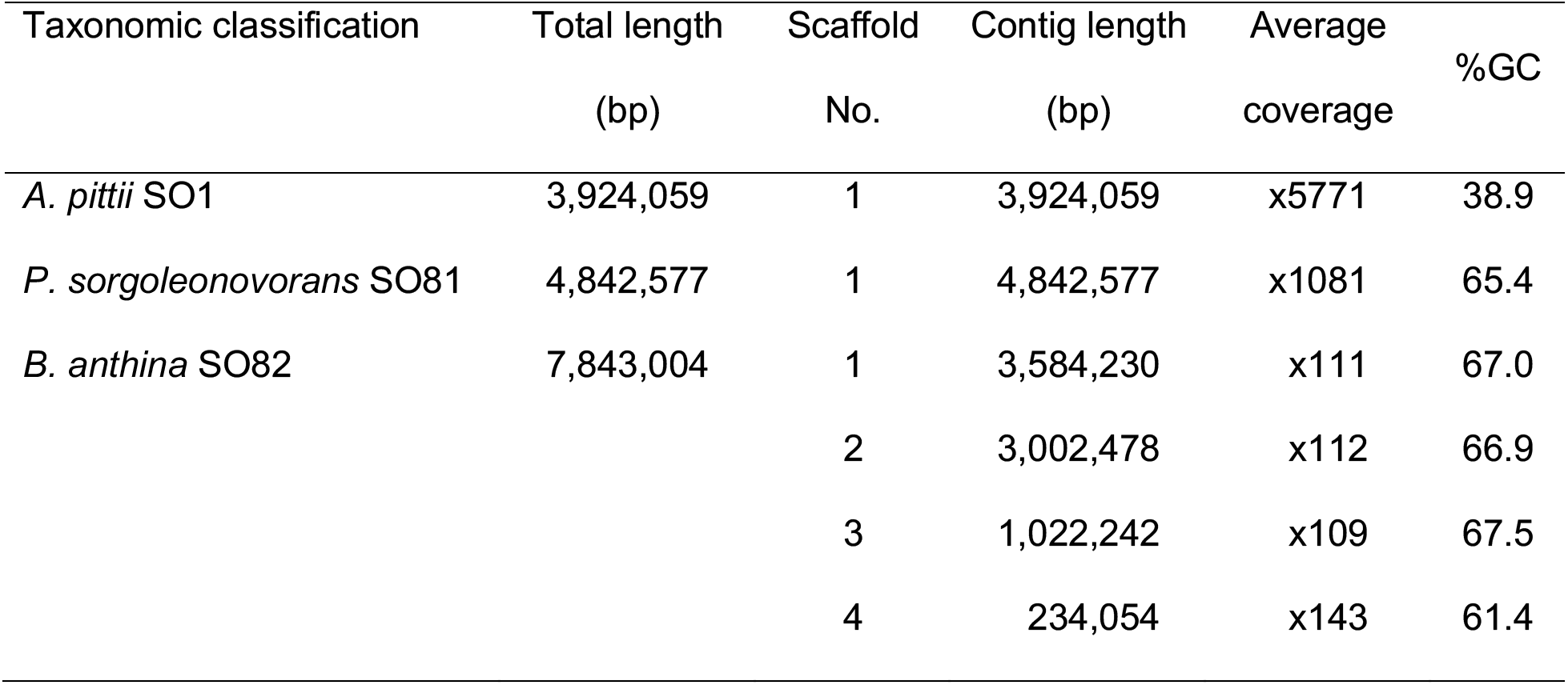
Statistics of whole genome sequencing of sorgoleone utilizing isolates

***SI Appendix*, Table S2a-S2c.** Transcriptome analysis of sorgoleone-grown strains (supplied as an Excel file)

***SI Appendix*, Table S3.** *P. sorgoleonovorans* SO81 RB-TnSeq analysis (supplied as an Excel file)

***SI Appendix*, Table S4.** Lookup table of the IMG and the GenBank locus tags (supplied as an Excel file)

**SI Appendix, Fig. S1.**
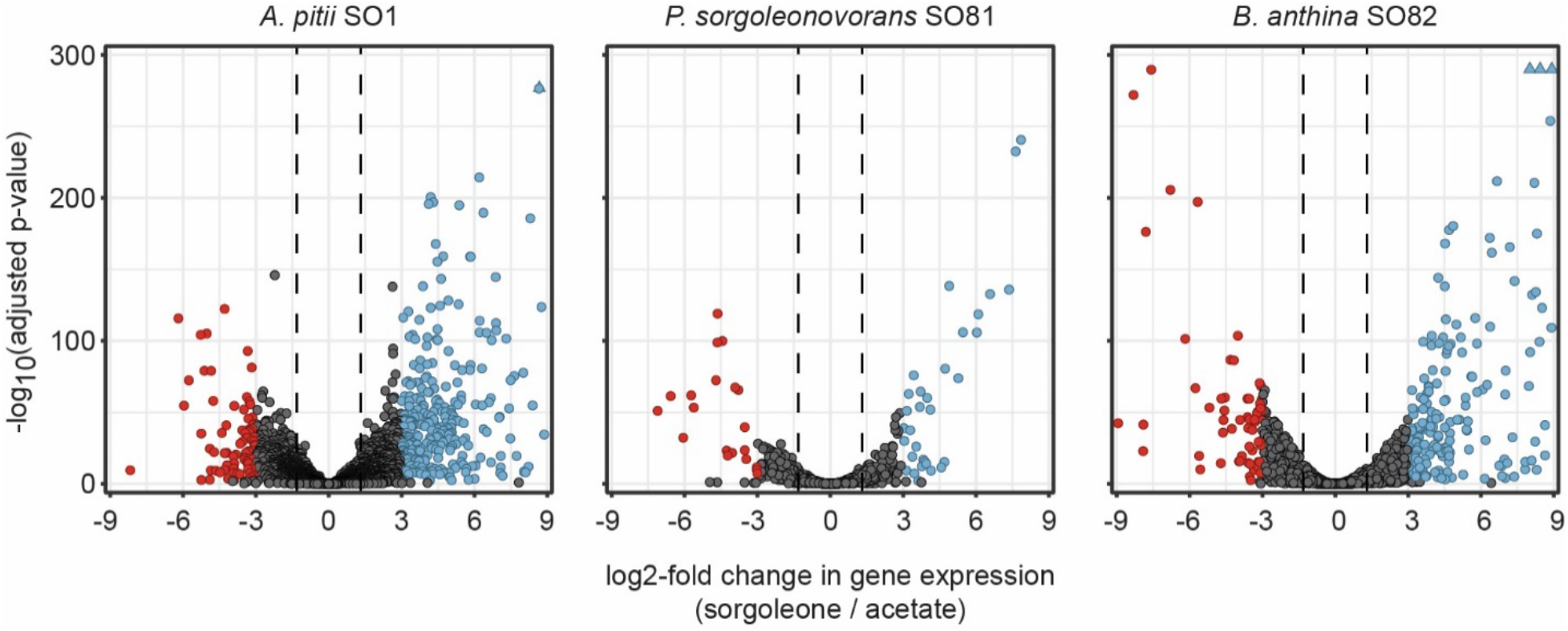
Summary of differential expression results. Differential expression of genes in *A. pitti* SO1, *P. sorgoleonovorans* SO81, and *B. anthina* SO82 during growth on sorgoleone versus acetate. Log_2_ -fold change in gene expression and -log_10_-transformed multiple test adjusted p-values are shown on the x- and y-axes, respectively. Positive numbers and negative numbers on the x-axis indicate higher expression during growth on sorgoleone and acetate, respectively. Dotted lines indicate ≥2.5-fold change in gene expression. Blue and red colored points indicate genes with both ≥8-fold change in expression and multiple test adjusted p-value <0.05 during growth on sorgoleone and acetate, respectively.

**SI Appendix, Fig. S2.**
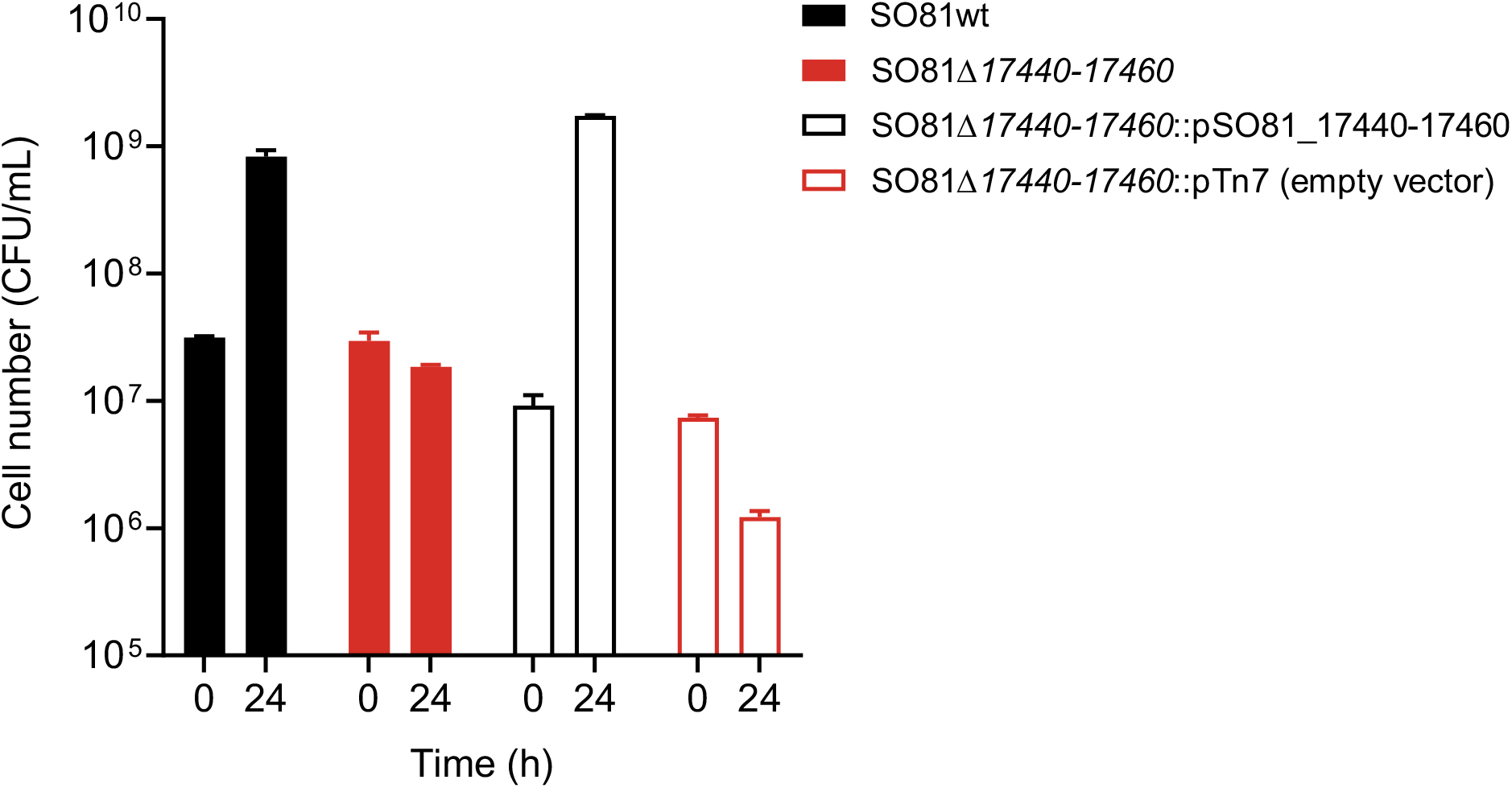
Growth *P. sorgoleonovorans* SO81 on sorgoleone depends on the srg genes SO81_17440-17460. Cells were incubated for 24h with 3 mM sorgoleone as a sole carbon source.

**SI Appendix, Fig. S3.**
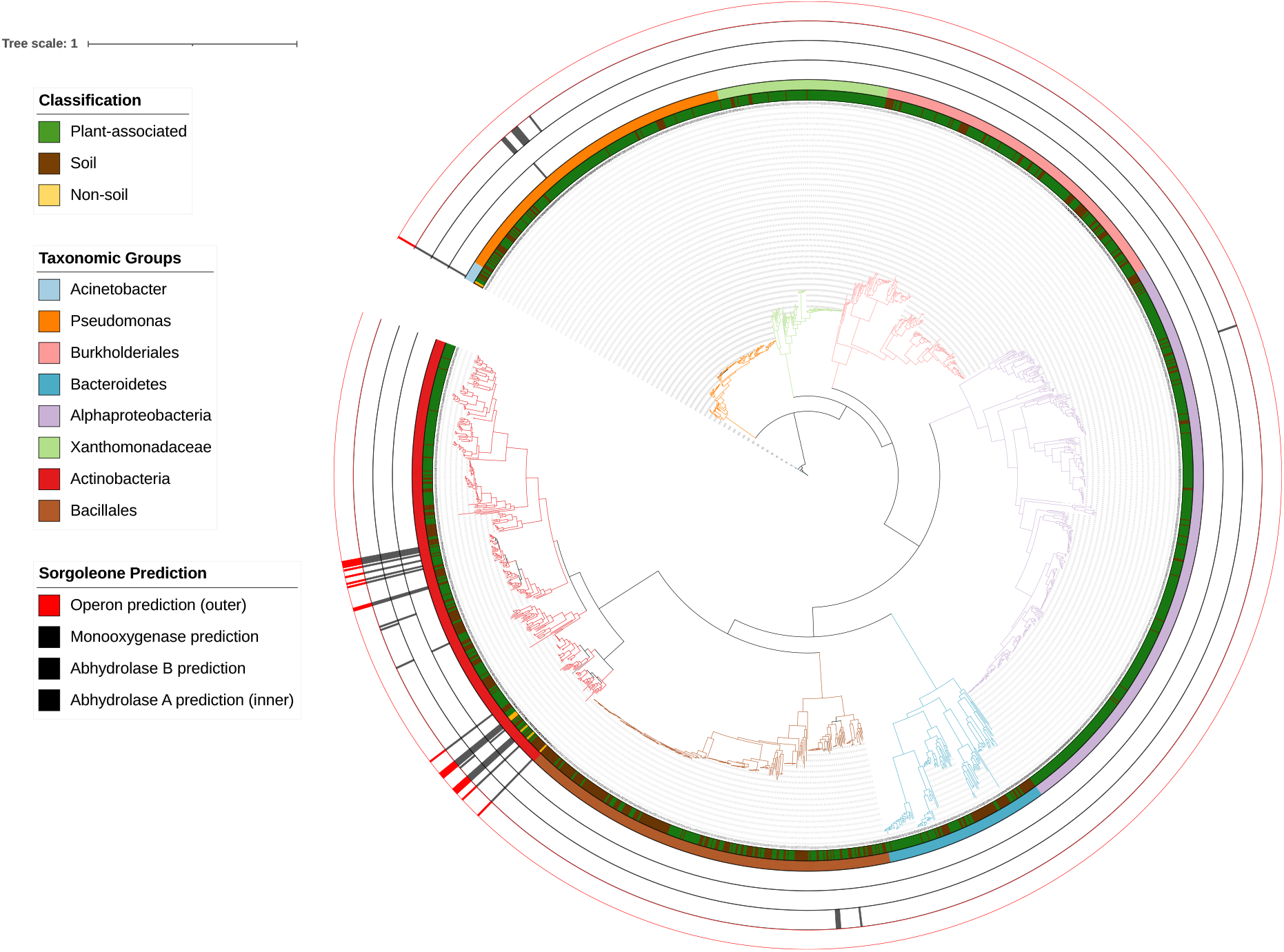
Prevalence of the *srg* cluster based on the Snekmer approach.

**SI Appendix, Fig. S4.**
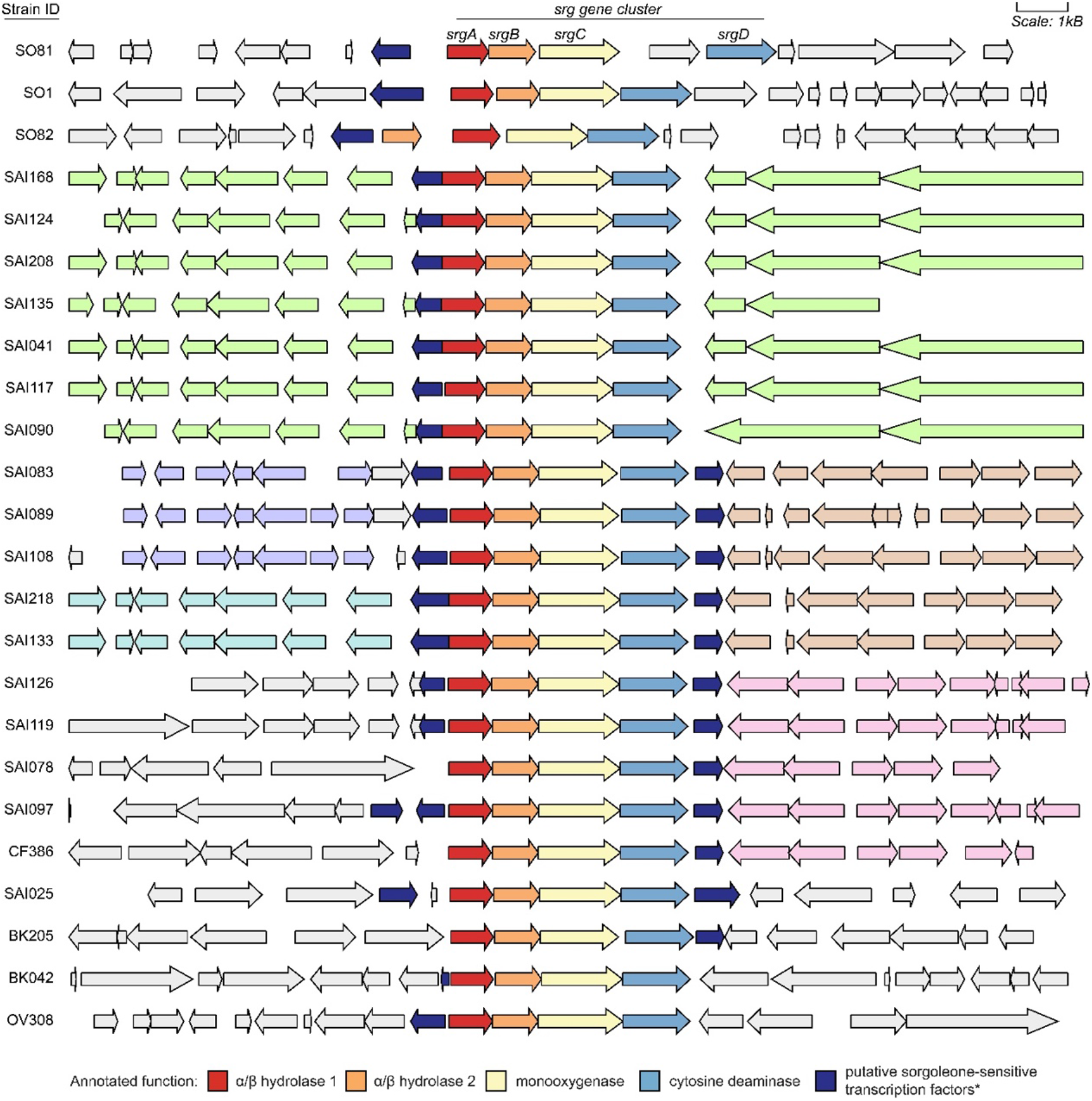
Alignment of *srg* gene clusters and neighboring genome sequences. Using the *srgC* genes identified by Snekmer analysis as central points, 20kb sequence regions were extracted from each genome containing a *srg* cluster (see color key for *srg* gene identifications). The proteins found within those regions were clustered using mmseqs2. Pastel colors of flanking genes indicate regions of conserved gene sequence and order across organisms. Genes with light gray fill did not cluster with any other genes in the analysis.

